# Modeling trophic dependencies and exchanges among insects’ bacterial symbionts in a host-simulated environment

**DOI:** 10.1101/086165

**Authors:** Itai Opatovsky, Diego Santos-Garcia, Tamar Lahav, Shani Ofaim, Laurence Mouton, Valérie Barbe, Einat Zchori-Fein, Shiri Freilich

## Abstract

Individual organisms are linked to their communities and ecosystems via metabolic activities. Metabolic exchanges and co-dependencies have long been suggested to have a pivotal role in determining community structure. Metabolic interactions with bacteria have been key drivers in the evolution of sap-feeding insects, enabling complementation of their deprived nutrition. The sap-feeding whitefly *Bemisia tabaci* (Hemiptera: Aleyrodidae) harbors an obligatory symbiotic bacterium, as well as varying combinations of facultative symbionts. We took advantage of the well-defined bacterial community in *B. tabaci* as a case study for a comprehensive and systematic survey of metabolic interactions within the bacterial community and their associations with documented frequency of bacterial combinations. We first reconstructed the metabolic networks of five common *B. tabaci* symbionts *(Portiera, Rickettsia, Hamiltonella, Cardinium* and *Wolbachia),* and then used network analysis approaches to predict: (1) species-specific metabolic capacities in a simulated bacteriocyte-like environment; (2) metabolic capacities of the corresponding species’ combinations, and (3) dependencies of each species on different media components.

The automatic-based predictions for metabolic capacities of the symbionts in the host environment were in general agreement with previously reported genome analyses, each focused on the single-species level. The analysis suggested several previously un-reported routes for complementary interactions. Highly abundant symbiont combinations were found to have the potential to produce a diverse set of complementary metabolites, in comparison to un-detected combinations. No clear association was detected between metabolic codependencies and co-occurrence patterns. The findings indicate a potential key role for metabolic exchanges as key determinants shaping community structure in this system.

## Importance

This study harnesses the rapid advances in tools developed within the newly emerging field of eco-systems biology to study a small, closed, well-defined micro-ecosystem of a bacterial community, allowing a detailed description of its trophic networks. In addition to indicating un-reported routes for complementary interactions between co-located symbionts of *Bemisia tabaci,* this study provides a generic tool for creating testable predictions of metabolic interactions in complex communities. Understanding the overall metabolic interactions in a given system is of key importance in ecology and evolution and can provide a powerful tool for expanding knowledge on inter-species bacterial interactions in various ecosystems.

## Introduction

Metabolic interactions are one of the main factors shaping communities and ecosystems by forming complex trophic networks. In bacterial communities, metabolic exchanges are ubiquitous and play a pivotal role in determining community structure (1–8). Bacteria also exchange metabolites with multicellular organisms, and such of mutualistic interactions have been a key driver of evolution, enabling eukaryotic expansion into new ecological niches and species diversification (9, 10). Among the most studied evolutionary radiations that has depended on symbiosis are the sap-feeding insects such as whiteflies, aphids, psyllids, cicadas and spittlebugs. All have intimate associations with maternally transmitted, intracellular bacteria that provide essential nutrients (mainly essential amino acids) and thereby enable dietary specialization on phloem or xylem sap of vascular plants (11–13) - a poor environment composed mainly of simple sugars and non-essential amino acids (14). The interaction with these inherited partners is obligatory for insect survival, and the bacteria are thus located inside specialized insect cells termed bacteriocytes. In addition, insects may harbor a diverse array of facultative, nonessential bacterial associates in the bacteriocytes or other body tissues (15). Facultative symbionts are suggested to serve as a “horizontal gene pool”, where variation in their combinations may have functional significance (16–19). Notably, since the obligatory symbionts are exposed to an irreversible process of genome reduction that can erode their metabolic potential (20), facultative symbionts can, in some cases, complement or replace parts of the lost functions (21–23).

In recent years, metabolic approaches, based on genome-driven network constructions, have been applied to predict the potential metabolic dependencies and metabolic exchanges between bacterial species (4, 8, 24). Newly developed tools for genome-based metabolic reconstruction enable predicting sets of interactions formed between species combinations, and the specific exchange of fluxes within multi-species systems (25, 26). Crossing such predictions with corresponding co-occurrence patterns allows deciphering the importance and meaning of variations in such bacterial assemblages (3, 27). To this end, multiple information layers are required, including symbiont co-occurrence patterns, environmental conditions, genetic background of both host and symbionts, and genome-driven predictions for symbionts’ potential activities. Here, based on the availability of both distribution patterns and bacterial genome sequences, we focused on exploring the functional significance of combinations of facultative symbionts in the sweetpotato whitefly *Bemisia tabaci* (Hemiptera: Aleyrodidae) and their potential role in shaping alternative community structures.

*Bemisia tabaci* is a major pest of several key crops worldwide (28) and is referred to as a complex of species, consisting of at least 28 morphologically indistinguishable, genetically delimited groups or species (29, 30). All whiteflies, including B. *tabaci,* harbor the primary symbiont “*Candidatus* Portiera aleyrodidarum” (hereafter *Potiera)* (31), which has undergone substantial genomic reduction as other obligatory symbionts (20), but is still able to produce most of the essential amino acids (32, 33). In addition, *B. tabaci* has been reported to harbor varying combinations of facultative symbionts, from bacterial genera *Rickettsia,* Hamiltonella, Wolbachia, Arsenophonus, Cardinium, Hemipteriphilus and Fritschea (34).

The occurrence and frequencies of combinations of these bacterial symbionts were investigated using a dataset of over 2,000 whiteflies, representing both the largest and the most comprehensive meta-study of insects for which communities of facultative symbionts have been described (34) MEAM1 and MED-Q1, the two most widespread genetic groups of *B. tabaci,* were found to typically harbor the facultative symbiont *“Ca.* Hamiltonella defensa” (hereafter *Hamiltonella)* in addition to the obligatory symbiont *Portiera.* A combination of *Hamiltonella* and “Ca. Rickettsia sp.” (hereafter *Rickettsia)* seemed to be unique to MEAM1 individuals, while combinations of *Hamiltonella* with either *“Ca.* Cardinium hertigii” or “Ca. Wolbachia sp.” (hereafter *Cardinium* and *Wolbachia* respectively) were unique to individuals of the MED-Q1 genetic group. Because the analysis revealed no correlation between specific facultative symbiont complexes and any of the environmental factors tested (34), we hypothesized that metabolic interactions may be involved in shaping the bacterial community structure. The recent release of the genome sequences of *Portiera, Rickettsia, Hamiltonella,* and *Cardinium* (23, 32, 35–40) has promoted analyses of interactions between the obligatory symbiont *Portiera* and its *B. tabaci*host (23, 33, 39, 41), the facultative symbionts and *B. tabaci* (23, 35, 36, 39), and the obligatory and facultative symbionts (23, 33). At both trophic levels, metabolic exchanges were suggested to be required for the completion of essential metabolic pathways. Branched Chain Amino Acids (BCAs), for example, are synthesized through *Portiera-host* complementary interaction (33, 39, 41) while lysine biosynthesis can occur via *Portiera-host* or *Portiera-Hamiltonella* complementation (23, 39).

As metabolic cross talk is suggested to convey functional capacities associated with specific species combinations, we conducted comparative-interaction analysis considering interactions formed between pairwise combinations of residing symbionts. We first reconstructed the metabolic networks of five symbionts *(Portiera, Rickettsia, Hamiltonella, Cardinium* and *Wolbachia),* and then used network analysis approaches to predict: (1) species-specific metabolic capacities in a simulated host’s bacteriocyte-like environment; (2) metabolic capacities of species’ combinations, and (3) the dependencies of each species on the different media components.

## Results

### Metabolic capacities of individual symbionts in the simulated bacteriocyte environment

The complete genomes of *Portiera, Cardinium, Hamiltonella,* and *Rickettsia* from *B. tabaci* MEAM1 and MED species were retrieved from public resources (Table 1) and the genome of *Wolbachia* was assembled *de novo* (Supplemental material, Table S1). All genomes were analyzed using a standard automated procedure followed by manual revision. For each bacterium, a metabolic-network was reconstructed based on the identification of its genome-derived enzyme content.

**Table 1:** Genomes list of obligatory and facultative symbionts of *Bemisia tabaci.*

| Symbiont | Host | Resource {GeneBank ID} (publication) | Number of ECs<sup>a</sup> |
| --- | --- | --- | --- |
| *Portiera* | MEAM1 | NCBI {CP003708.1} (40) | 103 |
| *Portiera* | MEAM1 | NCBI {CP003868.1} | 101 |
| *Portiera* | MED-Q1 | NCBI {CP003835.1} | 101 |
| *Portiera* | MED-Q2 | NCBI {CP003867.1} | 104 |
| *Cardinium* | MED-Q1 | NCBI {GCA_000689375.1} | 112 |
| *Hamiltonella* | MED-Q1 | NCBI {GCA_000258345.1} | 398 |
| *Rickettsia* sp. | MEAM1 | NCBI {GCA_000429565.1} | 247 |
| *Wolbachia* sp. | MED-Q2 | ENA {PRJEB15492} | 253 |

<sup>a</sup> Following annotation, filtering and manual curation. EC = enzyme commission.

Beyond the static representation of data as a network, computational simulations allow addressing the influence of environmental inputs (nutritional resources) on the network structure and composition, *i.e.,* the metabolic capacities of a species in a given environment, for example, in terms of its ability to produce essential metabolites. More specifically, expansion algorithms generate the set of all possible metabolites that can be produced given a set of starting compounds (source-metabolites) and a set of feasible reactions (42). We defined the starting compounds as a compilation of nutrients provided by the host whitefly in the bacteriocyte environment based on previous studies (33, 39, 41, 43). Our predicted bacteriocyte environment was composed of 50 compounds including ATP, co-factors and vitamins such as NAD+, heme and thiamine, six non-essential amino acids, and sugars (Table S2).

For each of the symbionts we simulated metabolic activity in the bacteriocyte environment and listed a sub-set of essential metabolites predicted to be produced (Table S3). It was found that most of the secondary symbionts are capable of producing nucleic acids (Fig. S1), whereas their ability to produce amino acids and co-factors varied (Fig. 1). *Portiera,* being an obligatory symbiont that has undergone substantial genomic reduction, was the most limited in its metabolic capacities. It was capable of synthesizing alanine and the essential amino acids threonine, methionine, tryptophan and phenylalanine (Fig. 1), in accordance with previous reports regarding its metabolic capacity and interaction with the whitefly host (32, 40). All of the facultative symbionts were capable of synthesizing the non-essential amino acid glycine, which was not produced by *Portiera.* As previously reported alanine is only produced by *Hamiltonella* and *Cardinium* (23, 35, 39). In addition, and in accordance with previous results, asparagine could be produced by the facultative symbionts *Hamiltonella, Wolbachia* and *Cardinium* (23, 39). Overall, the automatic-based predictions for metabolic capacities of the symbionts in the host environment generated by the model were in general agreement with previously reported genome analyses.

**Figure 1:**
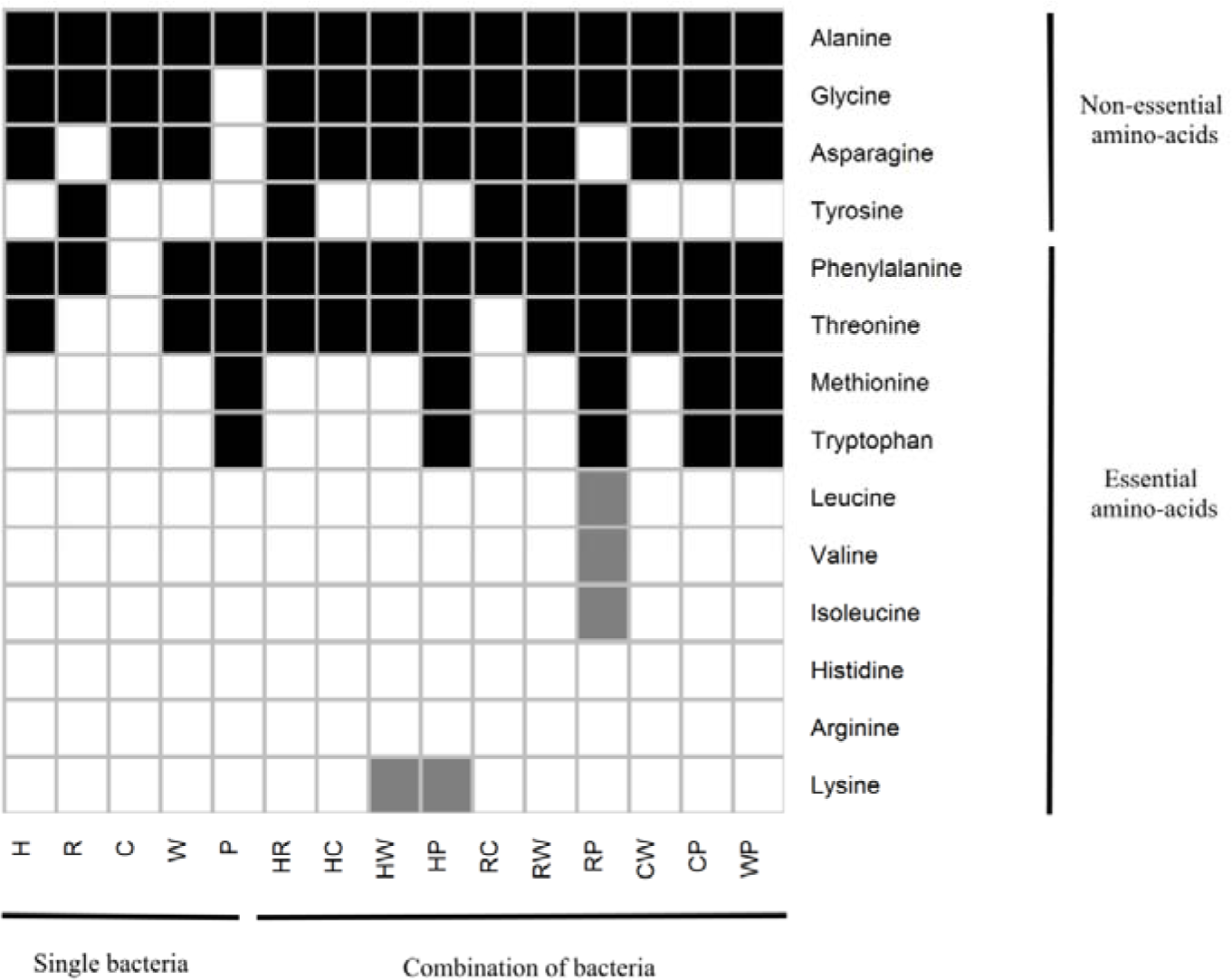
Predicted ability of single and pairwise species combinations to synthesize amino acids in the predicted bacteriocyte environment. Amino acids available in the bacteriocyte environment are not shown. Black/white/gray coloring of the cells - synthesis/no synthesis/production of complementary metabolites, respectively. P, C, H, R, W represent *Portiera, Cardinium, Hamiltonella, Rickettsia* and *Wolbachia,* respectively.

### Complementary production of amino acids

The genome-specific differences in the production of amino acids (Fig. 1) suggested that complementary metabolic interactions can potentially take place in the bacteriocyte ecosystem, increasing the total number of amino acids that can be synthesized by the residing bacteria. This is supported by a some established examples that demonstrate the coproduction of amino acids by bacterial combinations through complementation of metabolic pathways in various ecological systems, including insect-symbiont interactions (3, 44–46). To predict complementation patterns, we repeated co-growth simulations for pairwise combinations in the exact same environment as for single-species simulations. A metabolite was defined as “complementary’” if its synthesis requires a combination of bacterial species *(i.e.,* individual members of the combination cannot produce it). Overall, complementary interactions for the co-synthesis of four essential amino acids were detected (Fig. 1): lysine production by *Hamiltonella-Wolbachia* and *Portiera-Hamiltonella* combinations and production of the three BCAs (leucine, valine and isoleucine) by the *Portiera-Rickettsia* combination. While the complementation of *Hamiltonella-Wolbachia* for lysine production has not been previously reported, our results are in agreement with the possible cooperation of *Portiera and Hamiltonella* for its production (23, 39) The production of BCAs in the bacteriocyte environment has been suggested to take place through a complementary interaction between *Portiera* and *B. tabaci*. Our analysis suggested an alternative route for the production of BCAs through an interaction between the obligatory symbiont *Portiera* and the facultative symbiont *Rickettsia.* This previously unreported complementation is in agreement with identification of the ilvE gene in *Rickettsia* from *B. tabaci,* carrying the final reaction in the BCA-synthesis pathway (47).

### Profiles of complementary metabolites

Beyond the complementary production of amino acids, we recorded, for each pairwise bacterial combination, a vector describing the set of potential complementary metabolites (Table S4). The interactions formed between the most frequent symbionts - the obligatory symbiont *Portiera* and the partially fixated symbiont *Hamiltonella* - and the other symbionts, produced a high number of complementary metabolites per interaction (average of ~12; Table 2). In comparison, the lowest number of complementary metabolites was predicted for *Cardinium* (average of ~4, Table 2), the symbiont with the lowest number of appearances in the surveyed populations (34). Overall, the interaction matrix included seven occurring combinations (blue, Table 2) versus three non-occurring combinations (red), with an average number of ~12 versus ~3 complementary metabolites.

**Table 2:**
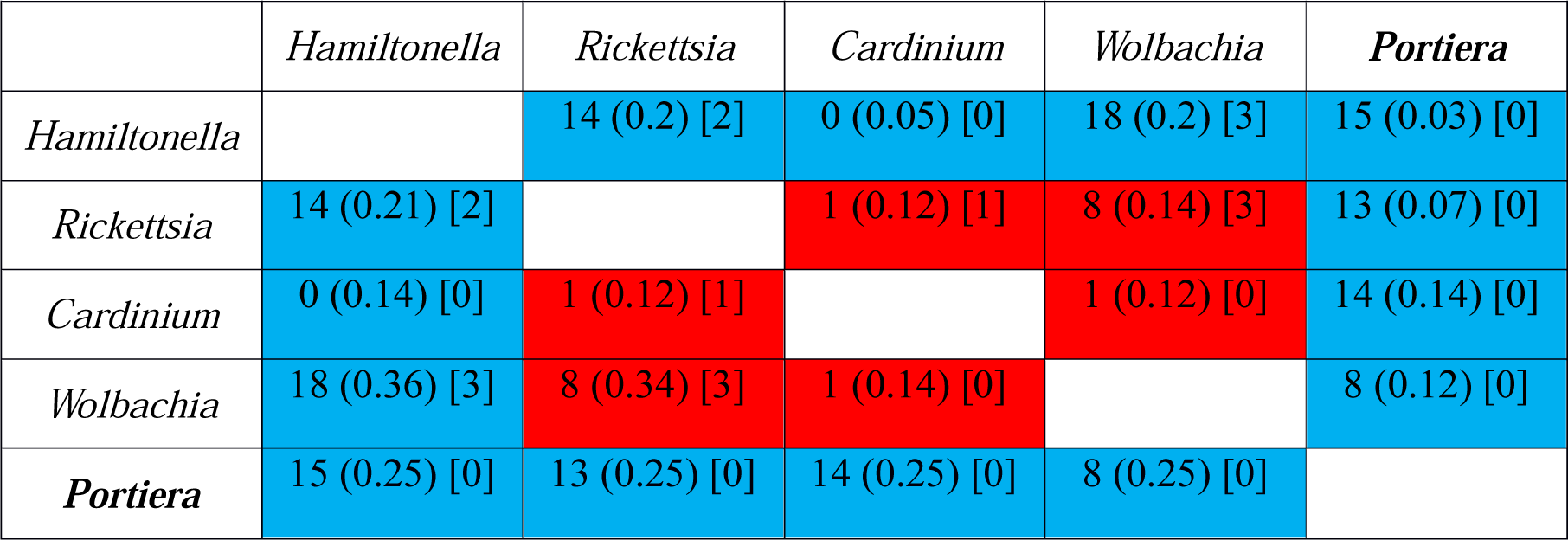
Predictions of pairwise interactions in the bacteriocyte system between occurring (blue) and non-occurring (red) pairwise combinations of symbionts. Occurrence versus nonoccurrence was determined according to a detailed survey of symbiont occurrence from 2030 whitefly individuals (34). The first value in each cell represents the number of complementary metabolites produced in each combination; the second value (in parentheses) represents the predictions of the competition values (Effective Metabolic Overlap); the third value (in square brackets) represents the number of source metabolites that induce codependency of both pair members. The primary endosymbiont is denoted in bold face.

Principle Component Analysis (PCA) of the complementary-metabolite vectors suggested four key types of interaction-groups (Fig. 2): *Portiera* associated interactions (with *Hamiltonella, Rickettsia* and *Wolbachia),* the two divergent Hamiltonella-associated interactions (with *Wolbachia* and *Rickettsia),* and the non-occurring combinations *Cardinium-Wolbachia,* and *Rickettsia-Wolbachia* and *Rickettsia-Cardinium* (red combinations in Table 2 and Fig. 2). *Cardinium*-*Portiera* combination is classified together with *Hamiltonella*-*Wolbachia* and not with the other *Portiera* associated combinations. Metabolites common to the Portiera-associated combinations included amino-acyl transferases and many primary metabolites such as amino acids and co-factors. Complementary metabolites common to the co-clustered *Portiera-Hamiltonella* and *Portiera-Wolbachia* combinations included potential precursors of methionine and purine/thiamine (Table S4); all potential interactions have been previously suggested for *Hamiltonella* (39), but not for *Wolbachia.*

**Figure 2:**
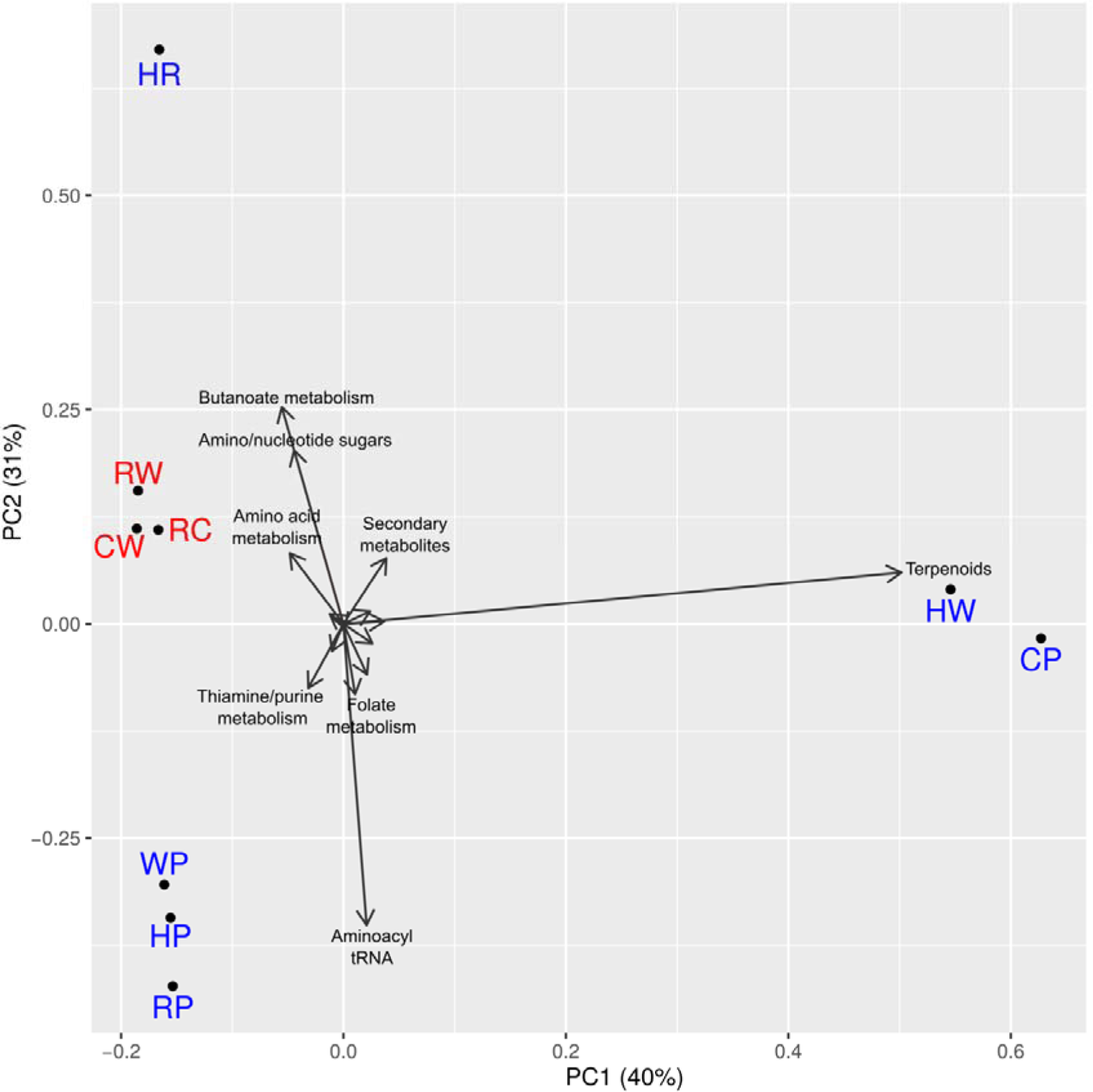
Principal Component Analysis (PCA) diagram of the synergistic metabolite profiles produced through pairwise interactions (Table S4). Synergistic metabolites are those whose synthesis requires the coexistence of both pair members and cannot be produced by either member alone in the predefined environment in which the simulations were carried out. Blue, co-occurring combinations; red, non-occurring combinations. P, C, H, R and W represent *Portiera, Cardinium, Hamiltonella, Rickettsia* and *Wolbachia,* respectively. HC combination has no synergistic metabolites and consequently is not represented. Vectors names represent the metabolic pathway of each synergistic metabolite in Table S4. For plotting reasons, only names of the most important vectors are displayed.

The relatively divergent clustering pattern recorded for the combinations of facultative symbionts *Hamiltonella-Wolbachia* and *Hamiltonella-Rickettsia* (Fig. 2) might be attributed to the fact that most of these metabolites are not common but rather interaction-specific: interactions between *Hamiltonella* and *Wolbachia* were mostly involved in the synthesis of secondary metabolites, mainly terpenoids; interactions between *Hamiltonella* and *Rickettsia* were mostly involved in butanoate and amino sugar metabolism (Table S4). Finally, non-occurring combinations typically led to a low number of potential complementary metabolites and were clustered.

### Co-dependencies of symbionts on specific media components

Under the assumption that highly similar metabolic demands may hint at resource competition and potentially lead to exclusion of the less fit competitor, the extent to which symbiont combinations rely on common resources was assessed. Scores were evaluated using NetCmpt, which provides predictions for the degree of effective metabolic overlap between pairs of bacterial species, ranging between 0 (no overlap) and 1 (complete overlap) (26). Scores are a-symmetrical whereas the effect of interactions on pair members is likely to differ *(i.e.,* one of the species is likely to be more affected than its potential competitor). The score is indicative of the effect of the column species over the row species. For example, *Hamiltonella* was almost unaffected by *Portiera* and *Cardinum* and was more sensitive to the presence of *Wolbachia* and *Rickettsia* (Table 2). Overall, pairwise scores were relatively low, ranging between 0.03 (the effect of *Portiera* on *Hamiltonella)* and ~0.35 (the effect of *Hamiltonella* on *Wolbachia* and *Rickettsia).* The observed average competition score, 0.18 (Table 2), was relatively low compared to an average of 0.36 calculated for other modeled bacterial communities (4). Notably, no significant difference was obsereved in the level of metabolic overlap between occurring versus non-occurring combinations (Table 2).

Since resource overlap is thought to determine community structure only under limited carrying capacity of the habitat (48), we further simulated species-specific growth in the bacteriocyte-like environment, rather than considering the generic optimal environment assumed by the NetCmpt tool. We estimated the specific qualitative effect of each metabolite on growth capacity following iterative removal of one component at a time. As expected, *Portiera* exhibited the most differentiated dependency profile of all symbionts (Fig. 3). In the specific bacteriocyte simulated environment, *Portiera* relied uniquely on D-ribose 5-phosphate, D-erythrose 4-phosphate and phosphoenolpyruvate for tryptophan production, as well as on L-homocysteine for methionine production. Metabolite dependencies that were common to more than a single symbiont included dependencies on the amino acids L-cysteine (*Wolbachia* and *Rickettsia)* and L-serine *(Cardinium, Hamiltonella* and *Wolbachia).* Hence, co-dependency might lead to a mutually exclusive distribution pattern, as suggested for *Wolbachia* and *Rickettsia* (34).

**Figure 3:**
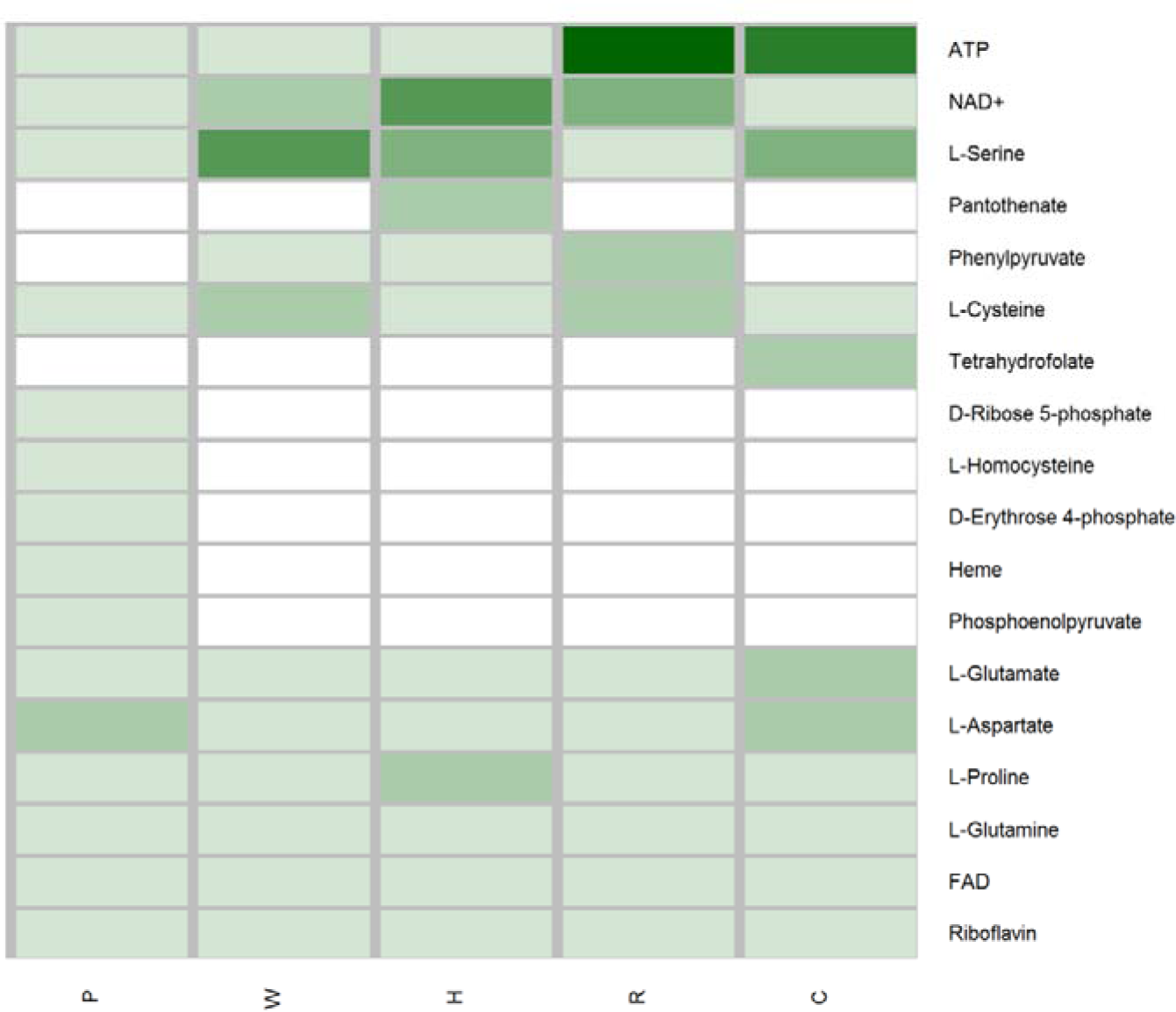
Reduction in symbiont’s ability to produce essential metabolites following removal of specific source metabolites (metabolites predicted to be available to the endosymbionts in the bacteriocyte). Only source metabolites whose removal affected at least one species are shown. P, C, H, R and W represent *Portiera, Cardinium, Hamiltonella, Rickettsia* and *Wolbachia*, respectively

In addition, common dependencies on NAD+ (*Hamiltonella, Wolbachia* and *Rickettsia)* and ATP (*Cardinium* and *Rickettsia)* reflected the energy production pathways of the corresponding symbionts. NAD+ dependent bacteria all have a citrate cycle requiring NAD+ as a reducing force. *Rickettsia* and *Cardinum,* both missing glycolytic pathways, rely on the host for ATP production. Though *Rickettsia* possesses a citrate-cycle, capable of producing ATP, its activation requires thiamine diphosphate, which was not present in our bacteriocyte environment. In our simulations, *Wolbachia* was the only symbiont that could produce thiamine diphosphate from the thiamine provided through the activity of thiamine diphosphokinase. Like *Cardinum, Portiera* does not possess either a citrate-cycle or glycolysis pathway. However, at least to a minimal amount, ATP production can potentially occur through the activity of ATP phosphoribosyltransferase in the histidine-metabolism pathway requiring D-ribose 5-phosphate as input. In addition, *Portiera* can also obtain ATP through carotenoid biosynthesis (49).

## Discussion

We harnessed the rapidly advancing tools developed within the newly emerging field of ecosystem biology to study a small, closed, well-defined micro ecosystem of a bacterial community. The focus on this unique community allowed exploring metabolic interactions between all relevant pairwise combinations, providing a detailed description of the trophic networks. Using simulation models to predict metabolic exchanges and co-dependencies we aimed to shed light on the role played by symbiotic interactions in shaping host ecology and how the ecology within the host can constrain community structure. The analysis was based on several assumptions and limitations that should be acknowledged: (1) we assumed a free flux of metabolites between the host and the symbionts and among the symbionts themselves. Several descriptions of the frequent exchanges in microbial communities support this assumptions (3, 50, 51). (2) The model is qualitative, only providing binary predictions for the production or absence of a metabolite rather than quantitative estimates for metabolite consumption/production as produced for stoichiometric networks using constraint based modeling. Hence, metabolites that are common resources for several symbionts might not induce competition, as they are not necessarily limiting. Similarly, the coproduction of nutrients might take place in negligible amounts, (3) the model is limited to the identification of metabolic interactions which are not likely to be the only factor affecting community structure. However, despite the inherent limitations of the approach, the analysis successfully captured previous genome-based predictions of metabolic complementations at host-symbiont and symbiont-symbiont levels in the bacteriocyte (23, 32, 39). Such evidence supports the relevance of our tool for the formulation of new, testable predictions of metabolic exchanges in an automated manner. Moreover, our simulations take into account a specific environment, hence reflecting the common notion that interactions are dynamic and can vary with the addition or depletion of nutrients (4, 6, 44).

This study focused on diet-limited insects that rely on obligatory associations with bacteria for complementation of their nutritional needs. The role of cooperative coevolution in selecting for traits that enable and stabilize such symbioses has been thoroughly discussed in the literature (10, 11, 13, 15, 16, 19, 52, 53). One of the most important negative ramifications of symbiotic alliances is the genome-reduction process in the obligatory symbionts that limits beneficial contributions (54). Consequently, a new symbiont may replace or supplement the capabilities of a previous one. The dynamic acquisition and loss of horizontally transmitted facultative symbionts enable the continuous persistence of many species. Although the facultative symbiont’s ability to colonize a new host is strongly influenced by metabolic similarities between the new and old host (35, 55), it also relies, at least to some extent, on the metabolic interactions that it forms with its new environment (56). Accordingly, transient bacterial species are expected to co-occur less frequently than expected by random chance if they are competing for limiting metabolic resources. Similarly, if their metabolic pathways are complementary with respect to the production of a mutually required resource, they are expected to co-occur more frequently than expected by random chance. Such interactions can also suggest a possible gain that compensates for the fitness cost of co-infections (56, 57).

Here, using automated tool rather than relying on genome-specific metabolic mappings (23, 35, 39), we predicted four previously un-reported routes for transient complementary interactions. These interactions can potentially increase the amount of the resulting amino acids in the bacteriocyte by providing alternative synthesis routes. Examples include complementation of the synthesis of BCAs is possible through the insect host *(B. tabaci)* obligatory symbiont *(Portiera)* interaction but also, by a previously un-reported interaction between *Portiera* and the facultative symbiont *Rickettsia.* Similarly, production of lysine as well as of the co-factor 5-methyl-tetrahydrofolate, the predominant form of dietary folate (58), occurs through the complementary *Portiera-Hamiltonella* interaction. The reported *Portiera-Hamiltonella* complementation of lysine could indicate a more intimate relationship between these symbionts, compromising the evolution of *Hamiltonella* toward a co-obligatory symbiont in some *B. tabaci* species (23, 36, 39). Though some of the complementary metabolites are redundant between co-existing interactions, they might suggest alternative production routes, possibly compensating for the limited transcriptional regulation of symbionts (59). Such complementation can be mutualistic, increasing the total amount of essential nutritional sources for all community members. Alternatively, it might only be beneficial for specific species and reflect a parasitic life style. For example, complementary production of BCAs is possible through *Portiera-Rickettsia* interactions. The *Rickettsia* from *B. tabaci* is part of the *R. bellii* group that includes many pathogenic members (60, 61). The complementation might reflect the dependency of *Rickettsia* on the BCA intermediates that it scavenges from the host-environment, bypassing the host’s control of BCA biosynthesis (47).

The model suggests several complementary pathways for metabolic co-production of additional metabolites, typical of *Portiera* interactions with the facultative symbionts. All of these interactions are involved in the production of metabolites compensating for the loss of aminoacyl-tRNAs in the *Portiera* lineage (L-tryptophanyl, N-formylmethionyl, L-methionyl and L-alanyl-tRNAs, Table S4) (33). Although these losses are assumed to reflect the dependency of *Portiera* on its host (30,31,58), the analysis suggests alternative routes for such complementation.

Complementary interactions also lead to the potential synthesis of secondary metabolites regulating host-parasitoid interactions(62, 63). For example, dimethylallyl diphosphate, a terpenoid, is involved in the metabolism of aphid’s alarm pheromones(64); sialic acids have diverse functions in host-bacteria interactions, including as signaling molecules and nutritional sources (65) (Table S4).

Specific combinations of co-occurring symbionts have been shown to correlate with delimited genetic groups of *B. tabaci* (34). Combinations of *Hamiltonella* with *Rickettsia* are unique to individuals from MEAM1, whereas combinations of *Hamiltonella* with *Wolbachia* are commonly found in individuals from MED-Q1. Notably, both combinations, which are highly dominant in their corresponding genetic group (34), have the potential to co-produce a diverse set of primary and secondary metabolites (14 and 18, respectively), which can increase host fitness, favoring their maintenance on this species. Unlike the relatively conserved profile of complementary metabolites produced through interactions between the obligatory and facultative symbionts, the complementary profiles formed by *Hamiltonella*-*Rickettsia* and *Hamiltonella-Wolbachia* are relatively diverse (Fig. 2), suggesting a biotype-specific functional adaptation. *Hamiltonella*-*Cardinum* combination is mainly found in the MED-Q1 group. This combination is less frequent (34), which could possibly be explained by their low complementation potential (zero metabolites). Consistent with these specific examples, we observed an overall trend of low complementary potential in non-occurring combinations in comparison to occurring ones. However, the limited sample size precludes significance of these observations.

While the analysis suggested an association between high-complementation and frequent cooccurrence no such indication was detected for competitive interactions (Table 2). One possible interpretation is that metabolic exchanges are more dominant in shaping bacterial communities (66). Indeed, whereas according to classical ecology theory, inter-species competition over common resources should lead to mutual-exclusion distribution patterns (48), relevant examples are rarely identified based on potential metabolic screens (3, 4, 67). A possibly explanation can be that only a narrow set of factors are quantitatively limited and therefore relevant for competition and determining community structure. To identify such potential limiting factors, we characterized metabolic co-dependencies between bacterial pairs. Predicted co-shared metabolites included the amino-acids L-cysteine (*Wolbachia* and *Rickettsia)* and L-serine *(Cardinium, Hamiltonella* and *Wolbachia).* Whereas *Hamiltonella-Cardinium* and *Hamiltonella-Wolbachia* combinations are frequent, *Wolbachia-Rickettsia* combinations are rare (34), indicating at cysteine as a potential limiting factor. Although cysteine is a non-essential amino acid that can be supplied by the host and is found in the phloem, it is the main sulfur source required for Fe-S protein biogenesis (68). In addition, common dependencies in NAD+ and ATP which reflect the energy-production pathways of the corresponding symbionts can have a strong influence on symbiont co-occurrences. For example, *Rickettsia* and *Cardinium,* both missing the glycolytic pathways and relying on their host for ATP production, are not found together in the host (34). In the *Rickettsia* genus, and other intracellular parasites, ADP/ATP translocases are known to play a crucial role in the exploitation of host ATP (60, 69). Interestingly, in *Cardinium* and related bacteria, ADP/ATP translocases are also present, indicating to a parasitic past (33, 70, 71). In contrast, it seems that *Wolbachia,* independent of its parasitic status, does not present (or has not acquired) the ADP/ATP translocases, relying on its own machinery to produce ATP (72).

Despite its obvious limitations, this model provides a tool for generating predictions for testable hypotheses of metabolic interactions in bacterial communities. Understanding the overall metabolic interactions in a given system is of key importance in ecology and evolution and can provide a powerful tool for expanding knowledge on inter-specific bacterial interactions in various ecosystems. With respect to applied aspects, symbiotic microorganisms have been shown to influence the success rates of various biological control programs of agricultural pests (73, 74). Attempts to establish more efficient pest-management strategies involve the removal of specific symbionts or the introduction of others, and our proposed model is expected to contribute to the efficiency and productivity of such efforts. The presented simple model system offers a level of tractability that is crucial for paving the way to the simulation, prediction and management of microbial communities that can expanded to more complex ecosystems, such as the guts of humans and livestock, water resources and soils.

## Materials and Methods

### Genome assembly and annotation

Relevant genomes were collected from multiple public sources (Table 1), with the exception of the *Wolbachia* genome which was assembled *de novo* using sequence data produced by a Genoscope-funded project (http://www.genoscope.cns.fr). The sequence was deposited in the European Nuclear Archive (http://www.ebi.ac.uk/ena/data/view/) under project number PRJEB15492. The procedure is fully described in the supplemental data.

A standard protocol for annotation retrieval was applied for all genomes. Annotations were carried out using several genome-annotation pipelines: IMG/M (75), Kbase (http://kbase.us/), Rast (76), MG-rast (76). To estimate the accuracy and comprehensiveness of the predictions, we benchmarked the EC (enzyme commission) predictions for the *Cardinium* genome, retrieved from the four pipelines, with annotations derived from a detailed manual curation. The IMG/G predictions were the most comprehensive and in highest agreement with the manual curation (Fig. S2). Hence, for consistency, annotations for all genomes were retrieved using the JGI platform. For *Portiera,* out of four published genomes (Table 1), annotations for CP003835.1 were considered in the analysis, based on cross-genome comparative analysis of the enzymatic sets and the annotation status (manually curated, Fig. S2). Following annotation retrieval from JGI, reciprocal BLAST searches were carried out to eliminate contaminated sequences between co-occurring symbionts. The phylogenetic origin of highly similar sequences was determined according to BLAST best hits.

Putative pseudogenes for all re-annotated genomes were predicted using GenePrimp (77). Manual inspection was performed for all candidate pseudogenes that had an assigned metabolic function (EC number). In addition, previous annotations of *Cardinium* and *Portiera* (32, 35) were used as supportive information for pseudogene cleaning in these species. Finally, predicted pseudogenes with valid EC accessions were removed from the predicted EC list before conducting follow-up analyses. The number of ECs annotated for each genome is indicated in Table 1. The final EC lists are provided in Table S5.

### Metabolic activity simulations

Metabolic activity simulations were carried using the Expansion algorithm (42) which allows predicting the active metabolic network (expanded) given a pre-defined set of substrates and reactions. The full expansion of the network reflects both the reaction repertoire of each species/species-combination and the primary set of compounds, termed here “source-metabolites”. Briefly, the algorithm starts with a set of one or more biochemical compounds acting as source metabolites for a feasible reaction, i.e., a reaction for which all required substrates are available. This reaction is selected out of the reaction pool and added to the network. In an iterative process, the products of the chosen reaction are turned into the new substrates, and so on. Processing of the starting-point compounds by relevant reactions increases the number of available compounds that can act as substrates for other, previously in-activated reactions. The network stops expanding when there are no more feasible reactions. Although, the closest organisms with a well-known and defined bacterciocyte environment are aphids, we decided not to use the information generated for this organisms, based on the long divergence time between aphids and whiteflies (more than 250 Mya) and differences in their symbiotic communities and their mode of transmission (53, 78-80). Here, we described the resources available in the whitefly bacteriocyte by compiling several such pre-published lists that are based on genomic-driven analyses of the whitefly genome (23, 32, 39, 41). The list is composed of metabolites produced by the host only, though each symbiont changes the environment by consuming/secreting unique set of metabolites. The limitation of the environment to host secreted metabolites allows predicting potential pairwise interactions that would otherwise be masked by alternative host-symbiont routes. These compounds were termed “source metabolites” (detailed in Table S2) and were used as starting points for unfolding a meta-network formed when considering all enzymes detected across all bacterial genomes, leading to the construction of niche-specific networks.

### Prediction of complementary interactions

Complementation was predicted through a three-stage model (1) constructing a combined set of metabolic reactions (EC accessions) for each pairwise combination; (2) simulating cogrowth of both individual and combined bacterial genera in the predicted environment; (3) comparing the set of metabolites produced by the combined genomes to those formed by the individual genomes. Complementary/Synergistic metabolites were those formed by species combinations but not by the individual species. A list of the complementary metabolites produced in each interaction and their mapping to KEGG pathways is provided in Table S4. PCA for the vectors of synergistic metabolites was carried out using R software (81).

### Prediction of co-dependencies in source metabolites

The competition scores for each pair of symbionts were calculated by the network-based tool NetCmpt (26). Beyond the quantitative estimates, NetCmpt was further extended to identify dependencies on specific source metabolites. To this end, growth simulations were carried in the bacteriocyte-like environment used throughout the analysis, rather than in the optimal environment used for the generic NetCmpt calculations. Within each simulation, the number of essential metabolites was determined (e.g., amino acids, nucleic acid and co-factors, Table S3) (26). Iterative simulations were carried out while removing one source metabolite at a time. For each iteration, the number of essential metabolites that could not be produced following the removal of a source metabolite was recorded. The procedure is illustrated in Fig. S3.

## Acknowledgments

We thank Genoscope (http://www.genoscope.cns.fr) for providing the RAW data for assembly of the *Wolbachia* genome.

